# HIPPIE: A Multimodal Deep Learning Model for Electrophysiological Classification of Neurons

**DOI:** 10.1101/2025.03.14.642461

**Authors:** Jesus Gonzalez-Ferrer, Julian Lehrer, Hunter E. Schweiger, Jinghui Geng, Sebastian Hernandez, Francisco Reyes, Jess L. Sevetson, Sofie R. Salama, Mircea Teodorescu, David Haussler, Mohammed A. Mostajo-Radji

## Abstract

Extracellular electrophysiological recordings present unique computational challenges for neuronal classification due to noise, technical variability, and batch effects across experimental systems. We introduce HIPPIE (High-dimensional Interpretation of Physiological Patterns In Extracellular recordings), a deep learning framework that combines self-supervised pretraining on unlabeled datasets with supervised fine-tuning to classify neurons from extracellular recordings. Using conditional convolutional joint autoencoders, HIPPIE learns robust, technology-adjusted representations of waveforms and spiking dynamics. This model can be applied to electrophysiological classification and clustering across diverse biological cultures and technologies. We validated HIPPIE on both *in vivo* mouse recordings and *in vitro* brain slices, where it demonstrated superior performance over other unsupervised methods in cell-type discrimination and aligned closely with anatomically defined classes. Its latent space organizes neurons along electrophysiological gradients, while enabling batch and individual corrected alignment of recordings across experiments. HIPPIE establishes a general framework for systematically decoding neuronal diversity in native and engineered systems.

## 1. Introduction

A thorough understanding of neuronal cell types, their locations within the brain, and how they connect is essential for decoding the functional properties of neuronal networks and their role in producing behaviors [1, 2]. Recent advances in high-throughput sequencing technologies have enabled researchers to gather extensive gene expression data from single cells, providing unprecedented insights into cellular diversity and function at a molecular level [3, 4]. While these transcriptional profiles provide valuable insights, they fail to capture essential functional characteristics, such as cellular morphology [5] and electrophysiology which are crucial for understanding the roles of cell types within their specific niches and provide physiological insight into cellular states [6].

To date, most cell-type classifications based on electrophysiology rely on low-throughput methods, such as whole-cell patch clamping [7]. However, several high-throughput technologies for electrophysiological characterization have emerged, including planar high-density microelectrode arrays (HD-MEAs) [8, 9, 10], flexible three-dimensional microelectrode array baskets [11], and high-density probes like Neuropixels [12, 13]. These technologies enable extracellular recordings at scale, allowing simultaneous monitoring of multiple units across different areas, thereby facilitating the study of network-level dynamics. High throughput electrophysiological recordings have traditionally distinguished broad cell-type categories using scalar measures, such as action potential width, to segregate putative excitatory “broad-spiking” neurons from “narrow-spiking” inhibitory neurons [14]. Furthermore, integrating these high-throughput approaches with genetic tools like optotagging [15, 16] provides a powerful way of identifying neuronal subtypes within these networks.

Efforts to classify extracellular electrophysiological data have diverged into supervised and unsupervised paradigms. Supervised methods like LOL-CAT [17], which maps spiking patterns to transcriptomic cell types, are constrained by sparse labeled datasets. Unsupervised approaches, such as nonlinear dimensionality reduction for clustering extracellular waveforms [18] or spiking patterns [19], leverage unlabeled data but operate on single modalities, neglecting complementary information. Multimodal integration has emerged as a priority, exemplified by PhysMAP [20], which adapts singlecell genomics strategies [21] to electrophysiology via UMAP and weighted nearest neighbors. However, UMAP’s inability to model sequential dependencies limits its efficacy for spike-time derived data. Alternative variational autoencoder (VAE) based frameworks [22] unify modalities through separate encoders but prioritize reconstruction over discrimination [23] and require spatial metadata absent in *in vitro* systems like organoids. Self-supervised learning offers a promising middle ground by generating supervisory signals from unlabeled data.

Advances in self-supervised learning, such as scGPT [24], highlight the potential of pretext tasks for representation learning, but domain-specific adaptations are needed. For instance, NEMO [25] employs contrastive learning to align average waveform and autocorrelogram embeddings, circumventing VAEs’ reconstruction bias. However, its assumption of modality-invariant neuronal signatures may not hold for dynamically coupled features (e.g., spike timing vs. waveform). Conditional VAEs (cVAEs) could address these limitations by balancing discriminative and generative objectives [26]. Unlike standard VAEs, cVAEs condition latent representations on auxiliary variables (e.g., experimental parameters), enabling targeted feature extraction [26]. In single-cell genomics, this strategy improves cell-type-specific generation [27], while in computer vision, it achieves attribute-controlled image synthesis [28]. Applied to electrophysiology, cVAEs could disentangle shared and modality-specific neuronal traits without spatial priors [26], making them adaptable to *in vitro* models. By explicitly modeling conditional dependencies (e.g., brain region or cell line), cVAEs may overcome the reconstruction-discrimination trade-off while integrating spike waveforms, temporal dynamics, and metadata into a unified latent space.

Here we present HIPPIE (High-dimensional Interpretation for Physiological Patterns in Intercellular Extracellular recordings), a cVAE framework designed for multimodal neuron classification and clustering by integrating extracellular action potential waveforms with spike-timing derived measurements. We benchmark HIPPIE against PhysMAP, the leading unsupervised method, demonstrating superior performance in cell-type identification and brain region classification across diverse recording technologies. HIPPIE achieves consistent gains in both inductive and transductive learning scenarios, highlighting its adaptability to real-world experimental constraints. By unifying feature extraction, clustering, and classification into a single framework, HIPPIE advances the interpretability and scalability of electrophysiological analysis.

## 2. Results

### 2.1. The HIPPIE pipeline

We developed HIPPIE, a deep learning framework that uses parallel cVAEs built from 1dResnet [29] encoder and decoder modules (Figure 1A-B). The HIPPIE pipeline takes as input spike sorted data from extracellular electrophysiology recordings. From the spike sorted data, the average waveform and interspike interval (ISI) distribution are calculated as 1d vectors, a categorical string representing the recording technology used in the experiment is also used as input to the network. Waveforms are averaged across spike times for each individual unit, and interspike interval distributions are computed from the spike train list using 1-millisecond bins, spanning a total of 100 milliseconds. The average spike waveform is calculated from a sample of 100 to 500 spikes, a range selected to balance accurate waveform representation with efficient memory usage. In our pipeline [30], we employed Kilosort2 for spike sorting. Kilosort2 uses template matching to compare individual spike waveforms, resulting in relatively low waveform variability across spikes. This approach allows us to obtain reliable spike representations even for low firing rate neurons with around 100 spikes. Given an average firing rate of 2 Hz, a 10-minute recording would yield approximately 1200 spikes, from which the 500-spike sample provides a representative subset. The length of the extracted waveform is 2.5 ms, which corresponds to 50 data points when sampled at 20 kHz. For higher sampling rates the number of points around the spikes can be adjusted to capture the same time window and then interpolated to 50 points to feed the algorithm. This time window captures the majority of the action potential duration in most neurons. If the algorithm were to be adapted for other electrophysiologically active cells, the time window should be adjusted accordingly.The preprocessing pipeline to obtain these processed features from raw voltage recordings is described in more detail in section 4.1. We focused on average waveform and ISI distribution because they have been shown to be some of the features that correlate the most with cell type and brain region [18, 31]. This framework could also be extended with additional measurements, such as autocorrelograms, layer-specific information, or other connectivity features. The outputs from the encoding layers of different modalities are integrated using a mixture module. In this work, the mixture module is implemented as a multilayer perceptron (MLP) [32], which takes the latent embeddings from both unimodal encoders and passes the output to the unimodal decoders. This setup enables the model to learn a shared representation through backpropagation.

**Figure 1:**
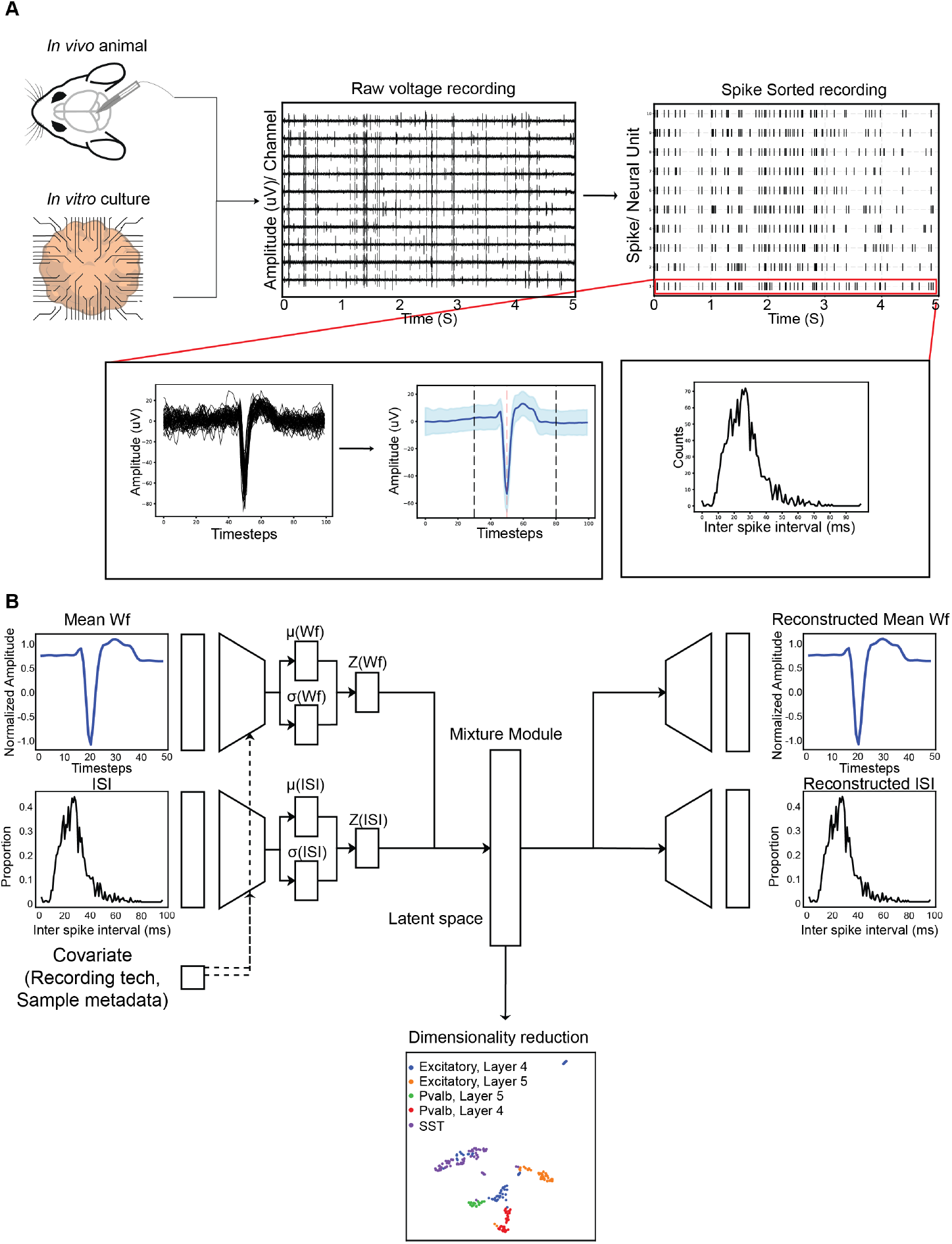
The HIPPIE pipeline. **A)**. Data from *in vivo* or *in vitro* experiments are spike sorted. From the reconstructed spike trains, all spike timings are collected for each unit. The interspike interval distribution is then computed based on the spike train data. To analyze waveforms, the spike timings are used to retrieve the corresponding raw voltage recordings, extracting a 2.5 ms window centered on the action potential peak. **B)**. 1 dimensional vectors representing the mean average waveform and interspike interval (ISI) distributions are fed int2o6paired conditional VAEs. The paired VAEs are trained in parallel, and a mixture module is employed to create a joint representation of the modalities in the embedding space, which will be used for downstream tasks after dimensionality reduction.

### 2.2 HIPPIE identifies anatomical locations better than other methods

To assess HIPPIE’s capability in generating meaningful brain regionspecific embeddings compared to the state-of-the-art method, PhysMAP [20], we conducted benchmark tests using datasets originally employed to validate PhysMAP’s ability to cluster transcriptomic cell types. However, our analysis prioritized the anatomical information within these datasets. Additionally, we supplemented these benchmarks with a new *in vitro* brain slice dataset generated by our group:

1. ***Juxtacellular Mouse S1 Area Dataset [15]*** : 293 units recorded in vivo from adult mouse primary somatosensory cortex (S1), spanning layers 2/3, 4, and 5 using custom tetrodes (Figure 2A).
2. ***CellExplorer Area Dataset [12]:*** 430 units from in vivo mouse hippocampal regions (CA1, CA3) and visual cortex (V1) recorded with Neuropixel 1.0 probes (Figure 2B).
3. ***Neonatal Mouse Brain Slice Dataset:*** This new dataset contains 2,220 units recorded *in vitro* from postnatal day 0–3 mouse brain slices (cerebral cortex, retrosplenial cortex, striatum, diencephalon) using Maxwell Biosystems’ MaxONE platform [33, 30] (Figure 2C).

**Figure 2:**
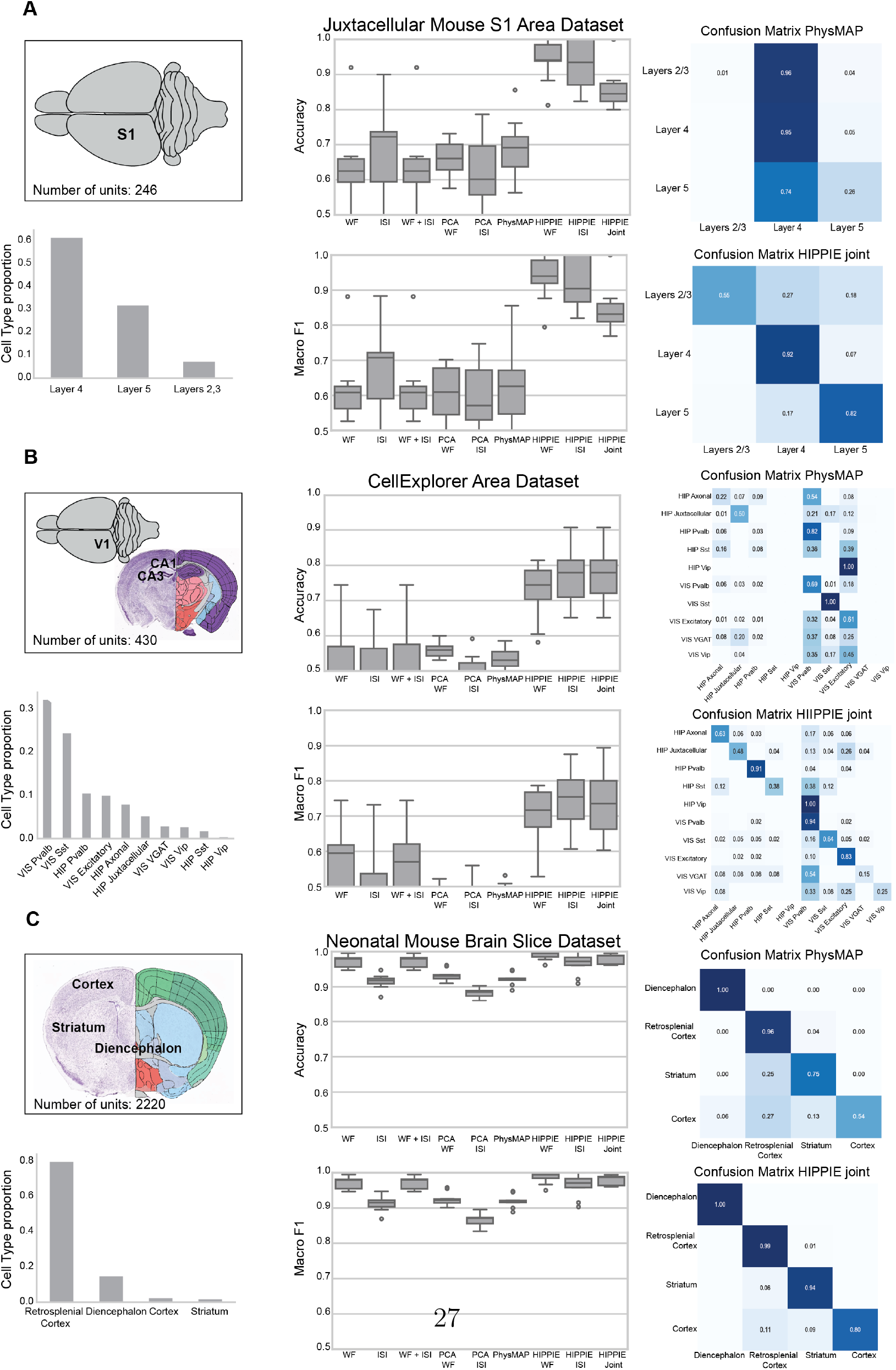
Anatomical locations benchmarks. Figure 2: **A)**. *Juxtacellular Mouse S1 Area Dataset Benchmark:* Left: Experimental location diagram (246 units) and neuronal layer proportions (L2/3, L4, L5). Middle: Boxplots showing Accuracy and Macro F1 across all tested dimensionality reduction methods. Right: Confusion matrices displaying layer-wise accuracy for PhysMAP (top) and HIPPIE joint modalities (bottom). **B)**. *CellExplorer Area Dataset Benchmark:* Left: Experimental locations (430 units) and neuronal subtype proportions (Pvalb, Sst, Vgat, Vip, Excitatory, Axonal, Juxtacellular) across Visual (VIS) and Hippocampal (HIP) areas. Middle: Boxplots summarizing Accuracy and Macro F1 for all tested dimensionality reduction methods. Right: Confusion matrices for PhysMAP (top) and HIPPIE joint modalities (bottom). **C)**. *Neonatal Mouse Brain Slice Dataset Benchmark:* Left: Experimental locations (2,220 units) and neuronal distribution across brain regions (retrospinal cortex, diencephalon, cortex, striatum). Middle: Boxplots presenting Accuracy and Macro F1 for all dimensionality reduction methods tested. Right: Confusion matrices showing accuracy for PhysMAP (top) and HIPPIE joint modalities (bottom). WF: Average waveform, ISI: Interspike Interval Distribution, Joint: Waveform and Interspike Interval

HIPPIE was pretrained using a leave-one-dataset-out strategy, incorporating the Allen OpenScope dataset [34], in-house recordings, and the above datasets. For each benchmark, the target dataset was excluded during pretraining and used only for evaluation. Following pretraining, models were fine-tuned on 80% of each target dataset (random split) and tested on the remaining 20%. Classification performance was evaluated using a K-nearest neighbors (KNN) classifier with 10-fold cross-validation across three input conditions: unimodal average waveform, unimodal ISI, and a combined multimodal input of average waveform and ISI (hereafter referred to as “joint”). HIPPIE outperformed other methods in predicting anatomical locations across all datasets, regardless of the deployed modality (Supplemental Tables 1–3, Figure 2), achieving the highest accuracy and macro-F1 scores. While PhysMAP [20] and PCA-based methods exhibited lower cross-validation variance compared to raw data analyses, their performance did not exceed that of unprocessed recordings. The UMAP representation of the embeddings obtained from the different methods also showcashes HIPPIE’s ability to create latent spaces with well diferentiated regions that show correspondence to different anatomical location(Supplemental Figure 1 A-C).

These results show that HIPPIE’s self-supervised architecture effectively extracts anatomical discriminative features from ISI distributions and average waveform, surpassing both PhysMAP and modality-agnostic dimensionality reduction approaches.

### 2.3 HIPPIE enables cross-sample generalization in anatomical classification

Conventional classifiers in electrophysiology often employ transductive splits, where neurons from all experimental samples are pooled before partitioning into training and testing sets [20, 25]. While this approach simplifies evaluation, it risks inflating performance metrics by allowing models to exploit sample-specific artifacts (e.g., electrode drift, session-specific noise) rather than biologically generalizable features [31]. In contrast, inductive splits enforce strict separation between training and testing samples, mimicking real-world scenarios where models must generalize to entirely unseen experimental preparations, a critical benchmark for robustness in neuroscience, where variability in recording conditions (e.g., electrode placement, animal-to-animal differences) remains a persistent challenge [31].

To rigorously assess HIPPIE’s generalizability, we analyzed the Allen Brain Institute’s Visual Coding dataset [34], the largest publicly available repository of *in vivo* Neuropixel 1.0 recordings from the mouse brain. Our analysis includes 16,564 units across 40 anatomically defined brain areas, collected from 47 adult mice. We evaluated HIPPIE under two distinct paradigms:

#### 1. Transductive Split

Using 80-20 stratified splits (balanced across brain areas), HIPPIE achieved near-perfect classification accuracy. The ISI modality outperformed other input types with 99.6% accuracy, closely followed by the waveform (98.4%) and joint (98.4%) modalities. In contrast, baseline methods, including PCA-based dimensionality reduction and PhysMAP [20] failed to exceed 15% accuracy (Supplemental Tables 1–3), indicating their reliance on dataset-specific confounders rather than invariant biological features.

#### 2. Inductive Split

To simulate real-world generalization, our splits consisted of 37 mice for training and 10 held-out animals for testing. HIP-PIE maintained robust performance, with the waveform (96.5%) and joint modalities (96.1%) modalities exhibiting minimal accuracy loss compared to transductive splits. The ISI modality experienced a sharper decline (99.6% to 91.4%). All baseline methods remained near chance (≤ 14.2 % accuracy), underscoring their inability to disentangle invariant neuronal properties from experimental noise (Figure 3).

**Figure 3:**
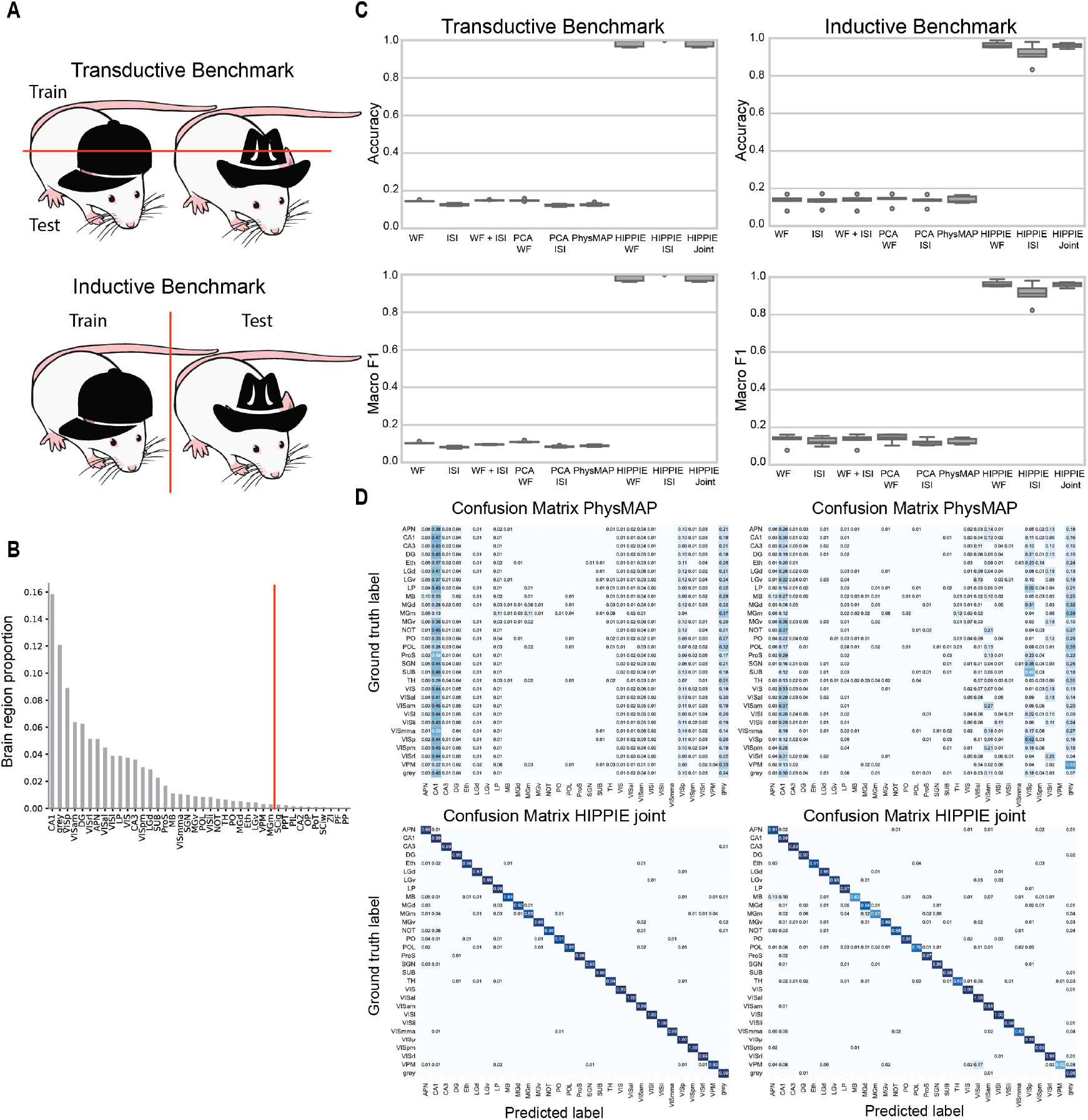
Inductive and transductive benchmarks for anatomical prediction. Figure 3: **A)**. Diagram illustrating the difference between transductive and inductive splits. **B)**. Proportion of cells per brain region in the dataset. Red line indicates the cells that were removed from the analysis due to having N *<* 50 **C)**. Accuracy and Macro-F1 boxplots comparing the performance of different dimensionality reduction methods using KNN probes. **D)**. Confusion matrices for transductive and inductive tests of PhysMAP and HIPPIE joint modality dimensionality reduction methods. WF: Average waveform, ISI: Interspike Interval Distribution, Joint: Average waveform and Interspike Interval Hippocampal structures include CA1, CA2, CA3, Dentate Gyrus (DG), Prosubiculum (ProS), and Subiculum (SUB). Thalamic structures include the Ethmoid Nucleus (Eth), Dorsal Lateral Geniculate (LGd), Lateral Posterior Nucleus (LP), Ventral Medial Geniculate (MGv), Posterior Complex (PO), Suprageniculate Nucleus (SGN), Ventral Posteromedial Nucleus (VPM), Parafascicular Nucleus (PF), Posterior Limiting Nucleus (POL), Ventral Lateral Geniculate (LGv), Posterior Intralaminar Thalamic Nucleus (PIL), Posterior Triangular Thalamic Nucleus (PoT), and the Medial Geniculate Complex, including its medial (MGm) and dorsal (MGd) parts. Visuocortical structures (VIS) include the Anterolateral (VISal), Anteromedial (VISam), Lateral (VISl), Primary (VISp), Posteromedial (VISpm), Rostrolateral (VISrl), and Laterointermediate (VISli) areas. Midbrain structures include the Anterior Pretectal Nucleus (APN), Superior Colliculus – Motor Related, Intermediate Gray Layer (SCig), Nucleus of the Optic Tract (NOT), Superior Colliculus – Motor Related, Intermediate White Layer (SCiw), Posterior Pretectal Nucleus (PPT), and Olivary Pretectal Nucleus (OP). The only hypothalamic structure included is the Zona Incerta (ZI). Cells labeled as grey represent units without an assigned brain region.

Visualization of HIPPIE’s latent representations revealed hierarchical clustering aligned with anatomical hierarchy even under inductive splits, confirming that its compressed embeddings encode biologically interpretable features rather than session-specific artifacts. These results establish HIPPIE as a model capable of generalizing across distinct experimental preparations while maintaining high anatomical resolution.

### 2.4 HIPPIE accurately classifies neuronal subtypes using waveform features

To assess HIPPIE’s ability to resolve cell-type-specific electrophysiological signatures, we analyzed three datasets combining *in vivo* extracellular recordings with optogenetic tagging of genetically defined neuronal subtypes. These datasets encompassed distinct recording technologies and brain regions:

#### 1. Juxtacellular Mouse S1 Cell Type Dataset (also known as Jianing dataset) [15]

This dataset comprised 224 units recorded juxtacellularly in the mouse primary somatosensory cortex using whole-cell patch pipettes. Cre-dependent ChR2 expression enabled optogenetic identification of Sst-positive and Pvalb-positive interneurons, with untagged putative excitatory neurons. Pvalb-positive interneurons and putative excitatory neurons are further subdivided by cortical layer (L4, L5).

#### 2. Extracellular Mouse Auditory Cortex (A1 Dataset) [35]

Recorded using NeuroNexus silicon probes, this dataset included 284 units from the auditory cortex. Pvalb-positive and Sst-positive interneurons were optotagged via Cre-mediated ChR2 expression, while putative excitatory neurons were identified by the absence of light response.

#### 3. CellExplorer Cell Type Dataset (Mouse Visual Cortex and Hippocampus) [12]

This Neuropixels 1.0-derived dataset contained 430 units spanning visual cortex and hippocampal regions. Vip-positive, Pvalbpositive, and Sst-positive interneurons were optotagged using subtype-specific Cre lines, with untagged neurons classified as putative excitatory.

For each dataset, HIPPIE was pretrained using a leave-one-dataset-out strategy, excluding the target dataset during pretraining. Following finetuning on 80% of the target data (stratified by cell type), performance was evaluated on the remaining 20% using KNN classification with 10-fold crossvalidation.

We found that HIPPIE’s waveform modality achieved the highest accuracy in cell-type discrimination across all datasets (Juxtacellular Mouse S1 96.42% Extracellular Mouse A1 Dataset 95.04% Cellexplorer Cell Type dataset 97.71%) (Figure 4; Supplemental Tables 4–6). The ISI and joint modalities exhibited slightly lower performance, though still superior to baseline methods (Juxtacellular Mouse S1 93.30% and 90.65%, Extracellular Mouse A1 Dataset 92.20% and 95.04%, Cellexplorer Cell Type dataset 94.13% and 93.64%). PhysMAP and PCA-based approaches demonstrated reduced cross-validation variance but failed to outperform analyses using raw data.Visual inspection of the embeddings produced by HIPPIE and PhysMap also reinforce the superior ability of HIPPIE to create meaningful latent spaces for cell type classification and clustering (Supplemental Figure 2 A-C).

**Figure 4:**
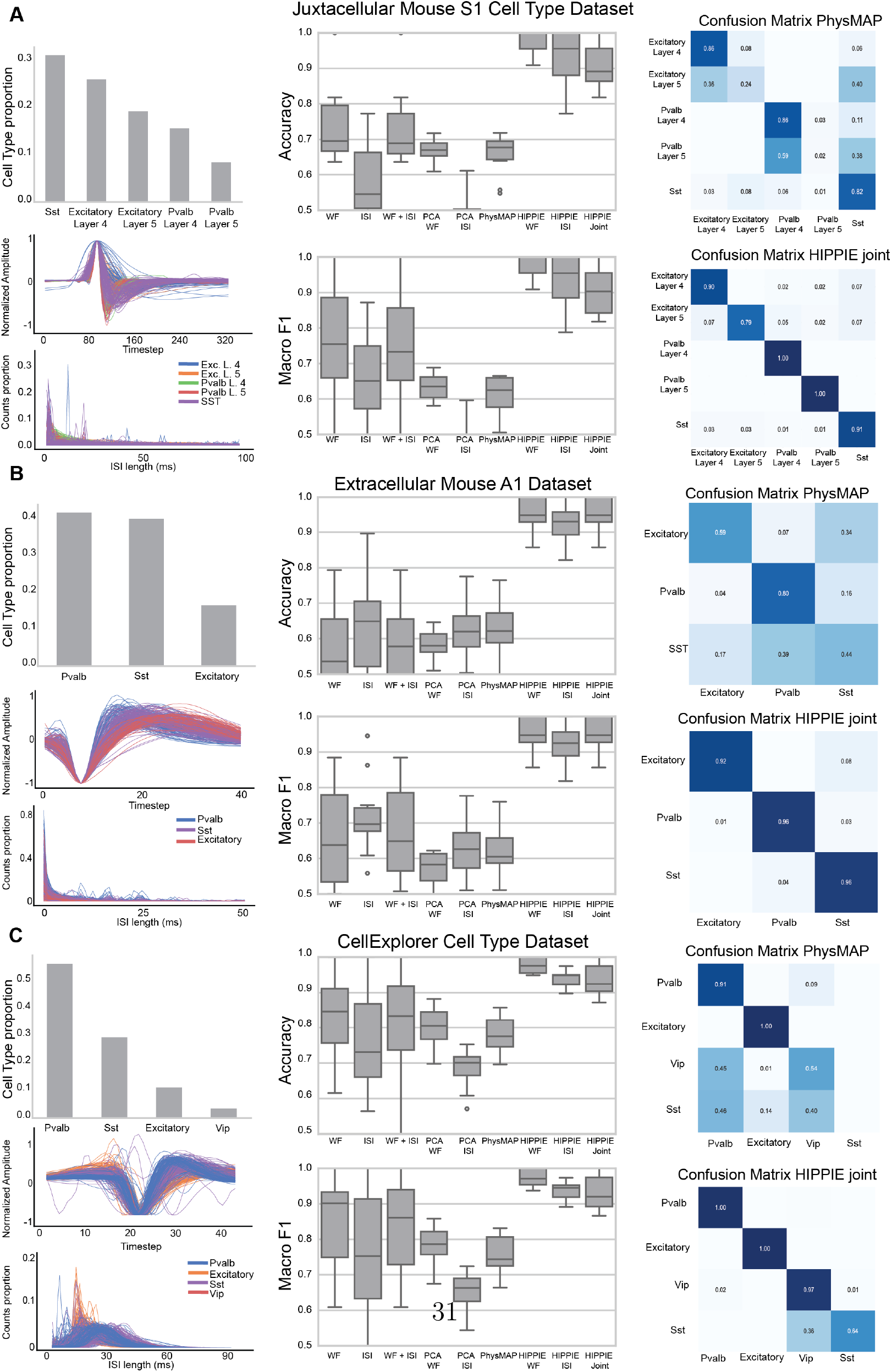
Cell types classification benchmarks. Figure 4: **A)**. *Juxtacellular Mouse S1 Area Dataset Benchmark:* Left: Neuronal subtype proportions (Sst, Pvalb Layer 4, Pvalb Layer 5, Excitatory Layer 4, Excitatory Layer 5) (top), amplitude (middle) and ISI distribution (bottom). Middle: Boxplots showing Accuracy and Macro F1 across all tested dimensionality reduction methods. Right: Confusion matrices displaying layer-wise accuracy for PhysMAP (top) and HIPPIE joint modalities (bottom). **B)**. *Extracellular Mouse Auditory Cortex dataset Benchmark:* Left: Neuronal subtype proportions (Sst, Pvalb Layer 4, Pvalb Layer 5, Excitatory Layer 4, Excitatory Layer 5) (top), amplitude (middle) and ISI distribution (bottom). Middle: Boxplots showing Accuracy and Macro F1 across all tested dimensionality reduction methods. Right: Confusion matrices displaying layer-wise accuracy for PhysMAP (top) and HIPPIE joint modalities. **C)**. *CellExplorer Cell Type Dataset Benchmark:* Left: Neuronal subtype proportions (Sst, Pvalb, Vip, Excitatory) (top), amplitude (middle) and ISI distribution (bottom). Middle: Boxplots showing Accuracy and Macro F1 across all tested dimensionality reduction methods. Right: Confusion matrices displaying layer-wise accuracy for PhysMAP (top) and HIPPIE joint modalities (bottom). WF: Average waveform, ISI: Interspike Interval Distribution, Joint: Waveform and Interspike Interval

These results establish HIPPIE as a robust framework for identifying neuronal subtypes in high throughout electrophysiology data. Furthermore, HIPPIE’s performance across juxtacellular, silicon probes, and Neuropixels 1.0 recordings highlights its generalizability to multiple experimental paradigms.

### 2.5. KNN probes are better suited for HIPPIE embeddings than Multilayer Perceptron probes

Although KNN probes were selected as the classification head of our model, alternative approaches, such as MLP probes, have also been employed to assess the discriminative capacity of the latent space in self-supervised models for high-throughput electrophysiological data [25]. To evaluate the effectiveness of our chosen approach, we compared the classification performance of KNN and MLP probes following HIPPIE dimensionality reduction. Across anatomical datasets, KNN consistently outperformed MLP for single-modality classification. In the CellExplorer Area Dataset [12], ISI embeddings classified with KNN achieved 76.98% accuracy, whereas MLP reached only 69.07%. The difference was even more pronounced for waveform embeddings, where KNN attained 72.79% accuracy compared to just 36.04% for MLP. However, for joint-modality classification, MLP outperformed KNN, achieving 90.23% accuracy versus 76.98%.

This trend was also evident in other anatomical datasets. For instance, in the Juxtacellular Mouse S1 Area Dataset, ISI embeddings classified with KNN achieved 93.08% accuracy, whereas MLP reached 91.16%. A similar pattern emerged with waveform embeddings, where KNN obtained 93.97% accuracy compared to MLP’s 72.31%. Yet again, for joint-modality classification, MLP surpassed both, achieving 94.20% accuracy compared to KNN’s 85.69%.

Notably, this trend did not hold in the Neonatal Mouse Slice ataset, where KNN outperformed MLP across all modalities. This substantial performance gap suggests that HIPPIE generates a feature space where local neighborhood relationships carry strong discriminative power. The superior performance of KNN implies that samples within the same class cluster more tightly in the HIPPIE-transformed space, making local distance-based methods more effective than the global decision boundaries learned by MLPs. However, for anatomical location classification, KNN’s non-weighted combinations of embeddings may limit its ability to effectively integrate information across multiple modalities.

In cell type classification tasks, where waveform features were the most informative, KNN consistently outperformed MLP. In the CellExplorer Cell Type Dataset [12], HIPPIE waveform classified with KNN achieved 97.7% accuracy, substantially surpassing the 81.40% obtained with MLP. This pattern held across additional datasets, including the Extracellular Mouse Auditory Cortex Dataset [35], where KNN reached 95.04% accuracy compared to 85.80% with MLP, and the Juxtacellular Mouse S1 Cell Type Dataset [15], where KNN achieved 96.42% versus MLP’s 71.60%.

Overall, these comparisons across ISI, waveform, and joint feature types indicate that HIPPIE’s dimensionality reduction generally produces a feature space where KNN outperforms MLP for single-modality classification, likely due to the strong local neighborhood structure in the transformed space. For joint-modality classification, MLP surpasses KNN in some anatomical classification tasks, but not all, while KNN consistently maintains an advantage in cell type classification. This suggests that MLP’s ability to learn weighted combinations of features provides an advantage for integrating multimodal anatomical information in specific cases, whereas KNN remains more effective when distinguishing cell types. A full set of results for MLP probes is available in Supplemental Table 7 and the boxplots showcashing accuracy and F1 can be found in Supplemental Figure 3.

## 3. Discussion

Autoencoders and related dimensionality reduction techniques have emerged as important tools for distilling high-dimensional neuronal recordings into interpretable latent representations. By compressing electrophysiological data while preserving biologically meaningful patterns, these methods enable the identification of cell-type-specific features and functional dynamics that are often obscured in raw recordings [22, 25]. The integration of multimodal neuronal data (such as extracellular waveforms and spike timing) into unified embedding spaces improves upon the limitations of single-modality analyses and enables a comprehensive neuronal characterization [20, 25]. Clustering algorithms applied to these latent spaces further refine our capacity to resolve neuronal diversity, uncovering non-linear relationships that may delineate novel cell subtypes or functional states.

Reproducibility, a persistent challenge in electrophysiology [36], can be tackled through the used of integration techniques and cross-sample alignment frameworks. This integration would not only enhance the transferability of analytical pipelines but could also empower validation of brain models, such as organoids, by enabling direct comparisons of neuronal phenotypes across studies. Advances in self-supervised learning are particularly promising for improving reproducibility, as these methods leverage high-throughput datasets (now widely accessible through repositories like DANDI) to learn generalizable representations of neuronal activity [30, 37], and tools such as HIPPIE and others have shown that conserved functional codes can be found across different animals [31]. By prioritizing invariant features across recordings, self-supervised models could overcome batch effects and technical variability inherent to electrophysiological experiments. The development of user-friendly graphical interfaces and web applications for implementing these models further enhances reproducibility by making complex analysis tools accessible to researchers without extensive programming expertise [38]. The democratization of computational methods reduces the technical barriers to adopt standardized analysis pipelines, enabling more laboratories to validate and build upon existing findings using consistent methodological approaches.

The rapid evolution of multimodal technologies is accelerating progress in neuroscience. Innovations such as *in situ* electro-sequencing [39], which integrates spatial transcriptomic profiling with electrophysiology, are generating rich datasets that link molecular identity to functional properties. Genetically enconded voltage indicators can also link molecular identity, electrophysiology, morphology and anatomical location [40]. Combined with next-generation brain observatories [41], these advancements will produce unprecedented volumes of neuronal data spanning brain regions, behaviors, and developmental stages. These large-scale datasets will enable the profiling of more cell types than ever before. However, they will initially be sparse, creating ideal conditions for self-supervised models like HIPPIE to excel. Initiatives such as the BRAIN Initiative Cell Census Network (BICCN) [1, 42] provide essential ground-truth cell-type atlases to validate machine learning-derived discoveries, while open-data platforms foster collaborative model development.

An algorithm for detecting brain regions and cell types from extracellular neural recordings also represents an important tool for the neurotech field. This tool could enhance brain-computer interfaces by identifying the brain region and cell types generating the signal, providing a better interpretation of user intent [43]. Brain region classification is particularly critical for understanding fundamental physiological differences between brain areas and for accurately targeting regions that are challenging to access via probe insertion. HIPPIE could provide estimates of probe location during experiments, significantly increasing the success rate of reaching target regions. This capability could be especially valuable in primates and human subjects, where histological information is often unavailable, and insertions rely on experimental heuristics that lack standardization across laboratories [44].

## 4. Methods

### 4.1 Data processing pipeline

For the dataset produced in house (Neonatal Mouse Brain Slice Dataset), raw recordings were processed following the pipeline described in [30]. The pipeline outputs a compressed ZIP file containing a spike data object structured as a NumPy array and a Python dictionary. This object includes the following components: a spike train list, a neuron data dictionary, the recording’s sample rate, and the electrode configuration. The neuron data dictionary provides spatial details, such as channel coordinates, neighboring channels, and spike features like waveforms and amplitudes. The indices of the spike train list correspond directly to those in the neuron data dictionary. Waveforms were averaged across spike times for each unit, and interspike interval distributions were computed from the spike train list using 1-millisecond bins, spanning a total of 100 milliseconds. The average spike waveform is calculated using a maximum number of 500 raw spikes. The length of the waveform is 2.5 ms, which is 50 data points with a 20 kHz sampling rate. Since HD-MEAs have a high number of electrodes and a small electrode pitch, one neuron’s signal can be recorded simultaneously by multiple electrodes. The neuron’s location on the HD-MEA is considered as the electrode with the maximum signal amplitude.

### 4.2 HIPPIE’s loss function

The cVAE loss function combines two essential components that work in concert to create a meaningful latent space. The first component is the reconstruction loss, which ensures the decoder can faithfully reconstruct the input data *x* from the latent representation *z*. The second component is the Kullback-Leibler (KL) divergence loss, which acts as a regularizer by encouraging the latent space representation to follow a prior distribution, typically a standard normal distribution. This combination is formalized as:

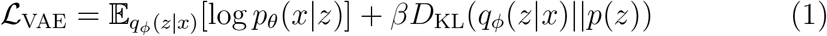

where *q*_*ϕ*_(*z*|*x*) represents the encoder network (also called the recognition model), *p*_*θ*_(*x z*) represents the decoder network (the generative model), and *p*(*z*) is the prior distribution. The hyperparameter *β* controls the trade-off between reconstruction quality and latent space regularity, with *β* = 1 recovering the standard VAE formulation.

When using a Gaussian encoder with diagonal covariance matrix, which is the most common choice, the KL divergence term has a convenient analytical form:

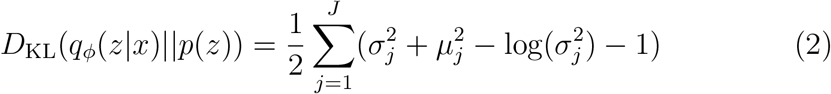

where *µ*_*j*_ and *σ*_*j*_ are the predicted mean and standard deviation for the *j*th dimension of the latent space.

This formulation ensures that the encoder learns to map inputs to a well-behaved latent space while maintaining the ability to reconstruct the original data accurately. Higher values of *β* place more emphasis on the regularization term, potentially leading to a more organized latent space at the cost of reconstruction fidelity, while lower values prioritize reconstruction quality.

### 4.5 Classification algorithms

The K-Nearest Neighbors (KNN) classification algorithm provides a robust framework for predictive modeling. K-nearest neighbors classification is done by majority vote of the labels of training samples closest to the test sample in feature space. We use euclidean distance for finding the nearest neighbors. Our approach can easily integrate a confidence scoring mechanism to evaluate prediction certainty, derived as the proportion of neighboring data points that share the majority class label, offering a probabilistic measure of classification certainty. By analyzing the label distribution among the k neighbors, the confidence score will range from 0 to 1, where higher scores reflect stronger consensus and better reliability.

MLP probing is used to evaluate the quality of VAE embeddings by training a supervised Multi-Layer Perceptron (MLP) classifier on the latent representations. The performance of this classifier indicates how well the learned embeddings capture task-relevant information. Compared to KNN probing, which assesses the structure of the latent space by measuring nearestneighbor classification accuracy, MLP probing provides a stronger test of linear separability and feature usefulness by optimizing a parametric model. While KNN reflects the local organization of embeddings, MLP probing evaluates their suitability for structured decision boundaries, making it a more flexible but potentially less interpretable evaluation method.

### 4.4. Evaluation Metrics

We employed KNN and MLP classifiers to assess the performance of dimensionality reduction techniques across neuronal cell type classification.

#### 4.4.1. Accuracy

Accuracy was calculated to evaluate model performance, defined as:

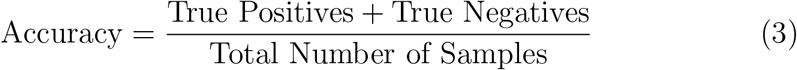

Where the total number of samples is the sum of True Positives, True Negatives, False Positives, and False Negatives.

#### 4.4.2. Macro F1 Score

The macro-averaged F1 score provides a comprehensive evaluation across all cell types:

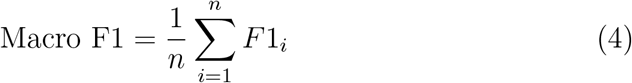

Where the F1 score for each class is:

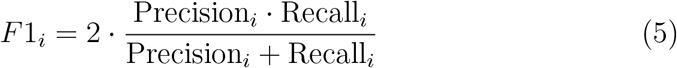

With Precision and Recall defined as:

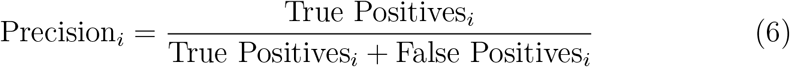

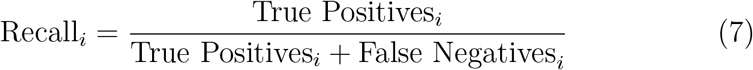

Performance metrics were estimated using stratified 10-fold cross-validation.

### 4.5. Code Library Details

The HIPPIE pipeline was designed with an easy to use application programming interface (API) to support a streamlined analysis with minimal code. To achieve this goal, the pipeline was constructed primarily using Py-Torch Lightning, a high-level library that aims to improve reproducibility, modularity, and simplicity in PyTorch deep learning code. We used Weights and Biases to visualize training metrics, including accuracy, F1 score, and loss, to facilitate the assessment of model performance.

All the code and instructions to use HIPPIE are available in the Braingeneers GitHub repository: https://github.com/braingeneers/Hippie

#### 4.5.1. Training details

For all experiments, models were trained on two phases: the pre-training phase and fine-tuning phase. For the pre-training phase, we train the model on every electrophysiology dataset, excluding whatever dataset we are measuring performance on. We perform a hyperparameter sweep over learning rate and the *β* parameter which determines the weight between the Kullback-Liebler loss of the latent space against a standard normal distribution and the *L*2 reconstruction error of the decoder. This is because *β* is a datasetdependent parameter. During pre-training, we incorporate the recording technology covariate but mask the class label embeddings to zero, even if labels may be available for some of the pre-training datasets. For fine-tuning and evaluation, we perform a K-fold cross-validation using 10 folds. For each fold, we train the model on the training set (80% of the data) and evaluate it on the remaining 20%. During fine-tuning, we use the class labels available to us and sample such that each mini-batch of training data contains an equal proportion of each label. This is to reduce the tendency of the model to predict only majority class labels while ignoring rare minority classes. We fine-tune with a learning rate equal to 1*/*10*× LR*, where *LR* is the pretraining learning rate. We fine tune for up to 50 epochs with a batch size of 128. For K-nearest neighbors evaluation, we fit the K-nearest neighbors algorithm to the training embeddings and then perform evaluation using the test embeddings compared with the test labels.

#### 4.5.2. Datasets

##### Juxtacellular Mouse S1 Dataset (Also known as the Jianing dataset)

The juxtacellular dataset collected by [45] was downloaded from the PhysMap’s github repository. Recordings were conducted in the primary somatosensory (barrel) cortex of genetically modified mice. Two transgenic lines were used: Pvalb-Cre::Ai32 to identify Pvalb-positive interneurons and Sst-Cre::Ai32 to identify Sst-positive interneurons.

These experiments focused on in vivo juxtasomatic electrophysiological recordings using glass micropipettes. Following recording, cells were filled with biocytin/neurobiotin, enabling morphological verification of cell types and alignment of recording depths with cortical layers. Spikes from ChR2-expressing neurons were identified based on laser-evoked responses to brief light pulses. To account for drift, spike waveforms were filtered. Depending on the experiment, we used either the cell type or the area label.

##### Extracellular Mouse Auditory Cortex Dataset (A1 Dataset)

The extracellular data analyzed from [35] was obtained from the PhysMAP GitHub repository. Pvalb-positive and Sst-positive interneurons were identified via opto-tagging in Pvalb-Cre::Ai32 or Sst-Cre::Ai32 mice. According to the dataset description, the identification of ChR2-expressing cells was performed conservatively, with positively identified cells defined as those exhibiting a significant increase in firing rate (p *<* 0.001) within the first 10 ms of stimulation onset.

Recordings were acquired from the primary auditory cortex using silicon probes, with optical stimulation delivered via a fiber positioned at the top of the probe recording sites. Spikes were sorted offline using Klustakwik, and only units with an ISI violation rate of less than 2% (spike intervals *<* 2 ms) per cluster were included in further analyses. The same parameters as those published in PhysMAP [20] were applied. Mean waveform for each unit were calculated using the channel with the highest spike amplitude.

##### CellExplorer Dataset (Mouse Visual Cortex and Hippocampus)

This dataset was downloaded from the PhysMAP github repository and includes recordings from the mouse primary visual cortex (V1), higher visual areas (HVAs), and hippocampus (CA1). Depending on the experiment, either anatomical region labels or cell type classifications were used.

Cell types were identified through opto-tagging in transgenic mouse lines. In the visual cortex, these included Pvalb-positive (Pvalb-IRES-Cre::Ai32), Sst-positive (Sst-IRES-Cre::Ai32), and Vip-positive (Vip-IRES-Cre::Ai32) interneurons. In the hippocampus, excitatory neurons and Pvalb-positive cells were identified, with V1 Pvalb-positive interneurons derived from Pvalb-IRES-Cre::Ai32 mice. Recordings were performed using Neuropixel 1.0 probes with optogenetic stimulation. [46, 47, 48]

##### Allen Institute’s Visual Coding Dataset

The Allen Institute’s Visual Coding dataset was accessed using Neurodata Without Borders (NWB) files retrieved via AllenSDK. Units were selected for further analysis based on the following default quality criteria: ISI violations *<* 0.5, amplitude cutoff *<* 0.1, and presence ratio *>* 0.9. To ensure robust statistical analysis, brain regions with fewer than 50 units were excluded. The spontaneous stimulus dataset also includes additional units from the “Functional Connectivity” set, complementing those from the “Brain Observatory 1.1” set.

#### 4.5.3. Neonatal Mouse Brain Slice Dataset

This dataset was generated specifically for this manuscript.

Postnatal day 0–3 mouse brains were extracted by a different group following ice-cold anesthetization prior to sacrifice. Immediately upon receipt, the tissue was sectioned into 300 µm slices using a Vibratome Leica VT1200S (amplitude: 1, advancement speed: 0.1 mm/s) in ice-cold sucrose solution (in mm): 72 sucrose, 83 NaCl, 2.5 KCl, 1 NaPO4, 3.3 MgSO4, 26.2 NaHCO3, 22 dextrose, 0.5 CaCl2. The slices were then transferred onto MaxWell Biosystems MaxOne HD-MEA chips.

Before experiments, the chips were pre-coated overnight at 37°C with 0.1 % Poly-L-ornithine (Millipore Sigma # P4957), followed by three washes with sterile water. Subsequently, they were coated overnight at 37°C with a mixture of 5 µg/mL Laminin (Millipore Sigma # L2020) and 1 µg/mL Fibronectin (Millipore Sigma DLW354008) in PBS.

The slices were maintained in BrainPhys Neuronal Medium (Stem Cell Technologies # 05790) supplemented with 10% (v/v) Hyclone characterized fetal bovine serum (Cytova # SH30071.03), 1X N-2 Supplement (Thermo Fisher Scientific # 17502048), 1X Chemically Defined Lipid Concentrate (Thermo Fisher Scientific # 11905031), 1X B-27 Supplement (Thermo Fisher Scientific # 17504044), 100 g/mL Primocin (Invivogen # ant-pm-05), and 1% (v/v) Matrigel Growth Factor Reduced (GFR) Basement Membrane Matrix (Corning # 354230). The slices were incubated at 37°C with 5% CO_2_.

For data processing, the data underwent the standard pipeline developed in the lab and published in the following manuscript [30]. We extracted the average spike waveform for each individual unit. Starting from raw recordings, these were spike sorted using Kilosort2, followed by auto-curation to get high-quality single-unit activity. Waveforms were then averaged across up to 500 spikes within a 5-ms window, using data from the channel capturing the neuron’s largest amplitude. All waveforms were centered at their peak, truncated to 1 ms before and 1.5 ms after the peak. For the ISI distribution, first, the timestamps of detected spikes were extracted for each unit. Then ISIs were computed as the differences between consecutive spike times and then they were binned into 100 equally spaced bins with 100 ms as the max threshold value.

### 4.6. Methods benchmarked against

1. **Raw data:** Raw data does not stand for the raw voltage recording obtained from MEAs but the processed mean waveform and ISI distribution for each unit as two 1-dimensional vectors. For the raw joint analysis we concatenated the two vectors into a larger 1-D array. We fed these vectors to the previously described KNN classification probe.
2. **PCA**: We applied PCA for dimensionality reduction on the normalized 1-dimensional vectors, retaining the first 10 principal components for the waveform and ISI modalities. We fed the reduced representation to the previously described KNN classification probe.
3. **PhysMAP:** PhysMAP is a method for identifying different cell types by integrating multiple physiological data modalities, such as neuronal waveform shape, ISI, peri-stimulus time histogram (PSTH), and derived electrophysiological metrics. It achieves this multi-modal integration by combining non-linear dimensionality reduction with a weightednearest neighbor (WNN) graph construction, inspired by Seurat v4 [21].

In this benchmark we only used it with the average waveform and ISI modalities. The code we used to run PhysMAP can be found in the following github repository https://github.com/EricKenjiLee/PhysMAP_Manuscript. We fed this reduced representation to the previously described KNN classification probe.

#### 4.6.1. Ethics statement

Experiments were performed according to protocols approved by the Institutional Animal Care and Use Committee at University of California Santa Cruz. Our group had access to tissue already sectioned provided by an independent lab.

## 5. Declarations

### 5.1. Author Contribution Statement

J.G.-F., J.L., and M.A.M.R. conceived the project. S.R.S., M.T., D.H., and M.A.M.-R. secured funding and supervised the work. J.G.-F., J.L., H.E.S., S.H., J.G., F.R., and J.L.S. performed the experiments. J.G.-F. J.L., and M.A.M-R. wrote the manuscript with contributions from all authors.

### 5.2. Lead contact

Further information and requests for resources and code should be directed and will be fulfilled by the lead contact, Mohammed A. Mostajo-Radji.

### 5.3. Material availability

No new materials were produced for this study.

### 5.4. Data availability

The majority of data used for this study was downloaded from existing datasets. The Neonatal Mouse Brain Slice dataset can be obtained upon request to the corresponding author.

### 5.5. Code availability

All code for HIPPIE has been deposited in https://github.com/braingeneers/Hippie

## 5.7. Acknowledgments

We would like to thank Aidan Schneider for his valuable feedback on this manuscript. In addition, we would like to thank Tommy Finn for providing mouse tissue for this project. This work was supported by Schmidt Futures (SF857) to S.R.S., M.T. and D.H.; National Human Genome Research Institute (1RM1HG011543) to S.R.S., M.T. and D.H.; National Science Foundation (NSF) (NSF2134955) to S.R.S., M.T. and D.H., (NSF2034037) to M.T.; the National Institute of Mental Health (1U24MH132628) to D.H. and M.A.M.-R.; the California Institute for Regenerative Medicine (CIRM) (DISC4-16285) to S.R.S., M.T. and M.A.M.-R., (DISC4-16337) to M.A.M.-R.; the University of California Office of the President (M25PR9045) to S.R.S., M.T., and M.A.M-R. H.E.S. was partially supported through the Graduate Research Fellowship Program (GRFP) of the NSF. S.H. was partially supported by the UC Doctoral Diversity Initiative (DDI-UCSC-IBSC) program. F.R. was supported by the CIRM Bridges to Stem Cell Research program awarded to Berkeley City College. We are thankful to the Pacific Research Platform, supported by the National Science Foundation under Award Numbers CNS-1730158, ACI-1540112, ACI-1541349, OAC-1826967, the University of California Office of the President, and the University of California San Diego’s California Institute for Telecommunications and Information Technology/Qualcomm Institute.

