## Supplemental Figures 1-3 and Supplemental Tables 1-7 for "HIPPIE: A Multimodal Deep Learning Model for Electrophysiological Classification of Neurons"

Supplemental Figure 1: UMAP plots for the different dimensionality reduction methods applied to the benchmarks for anatomical location prediction
Supplemental Figure 2: UMAP plots for the different dimensionality reduction methods applied to the benchmarks for cell type prediction
Supplemental Figure 3: K-Nearest Neighbors swap with Multilayer perceptron probe (MLP).

Supplemental Table 1: Juxtacellular Mouse S1 Area Dataset benchmark results.
Supplemental Table 2: CellExplorer Area Dataset benchmark results
Supplemental Table 3: Mouse slice P0 and P3 benchmark results
Supplemental Table 4: Juxtacellular Mouse S1 Cell Type Dataset benchmark results
Supplemental Table 5: Extracellular Mouse A1 Dataset benchmark results
Supplemental Table 6: CellExplorer Cell Type Dataset benchmark results
Supplemental Table 7: MLP probe results for all datasets

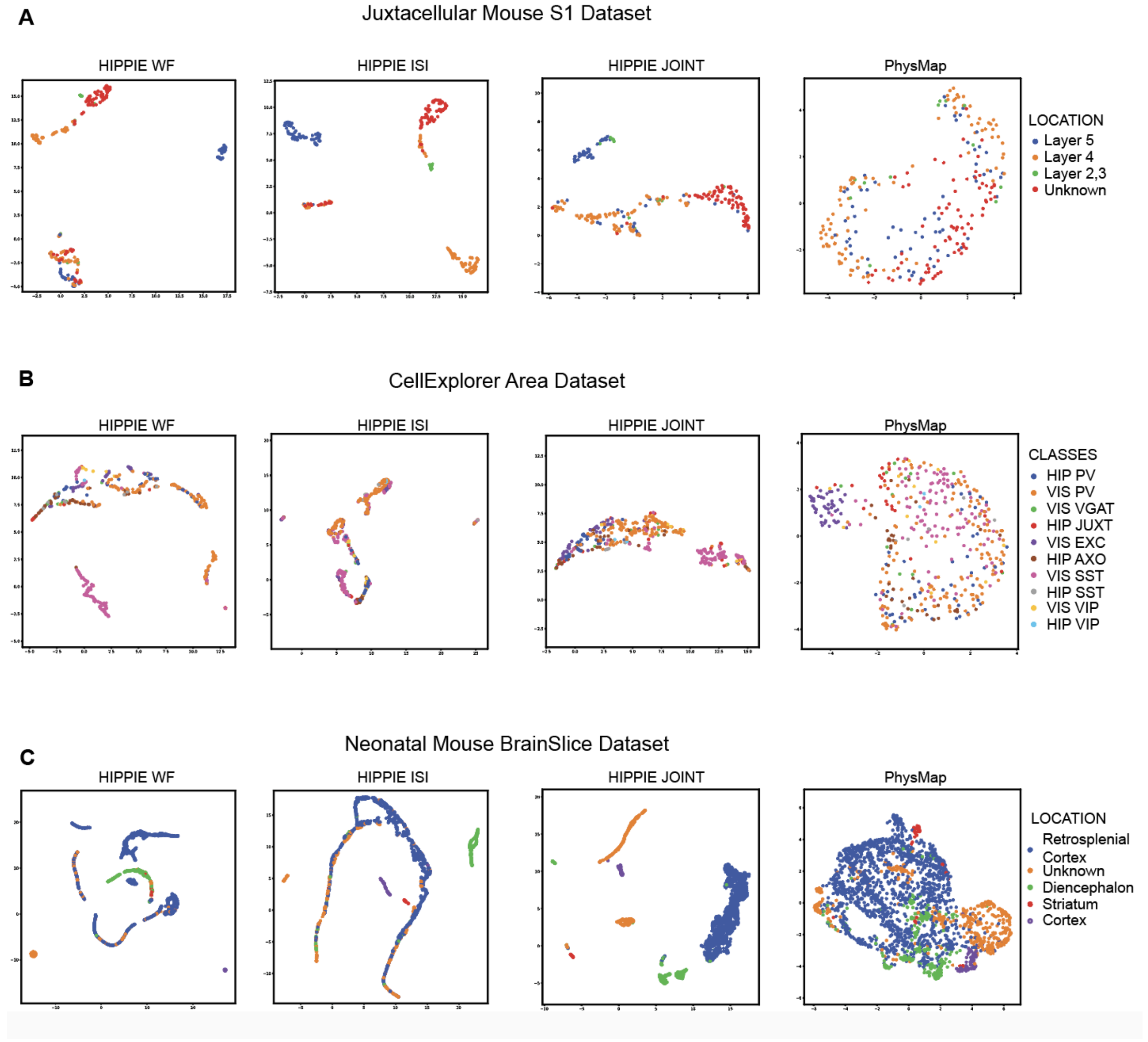


**Supplemental Figure 1**: **UMAP plots for the different dimensionality reduction methods applied to the benchmarks for anatomical location prediction.** A) UMAP representations for the latent space obtained from HIPPIE (Average Waveform, ISI, Joint) and PhysMap for the Juxtacellular Mouse S1 dataset. B) UMAP representations for the latent space obtained from HIPPIE (Average Waveform, ISI, Joint) and PhysMap for the CellExplorer Area dataset. C) UMAP representations for the latent space obtained from HIPPIE (Average Waveform, ISI, Joint) and PhysMap for the Neonatal Mouse BrainSlice dataset.

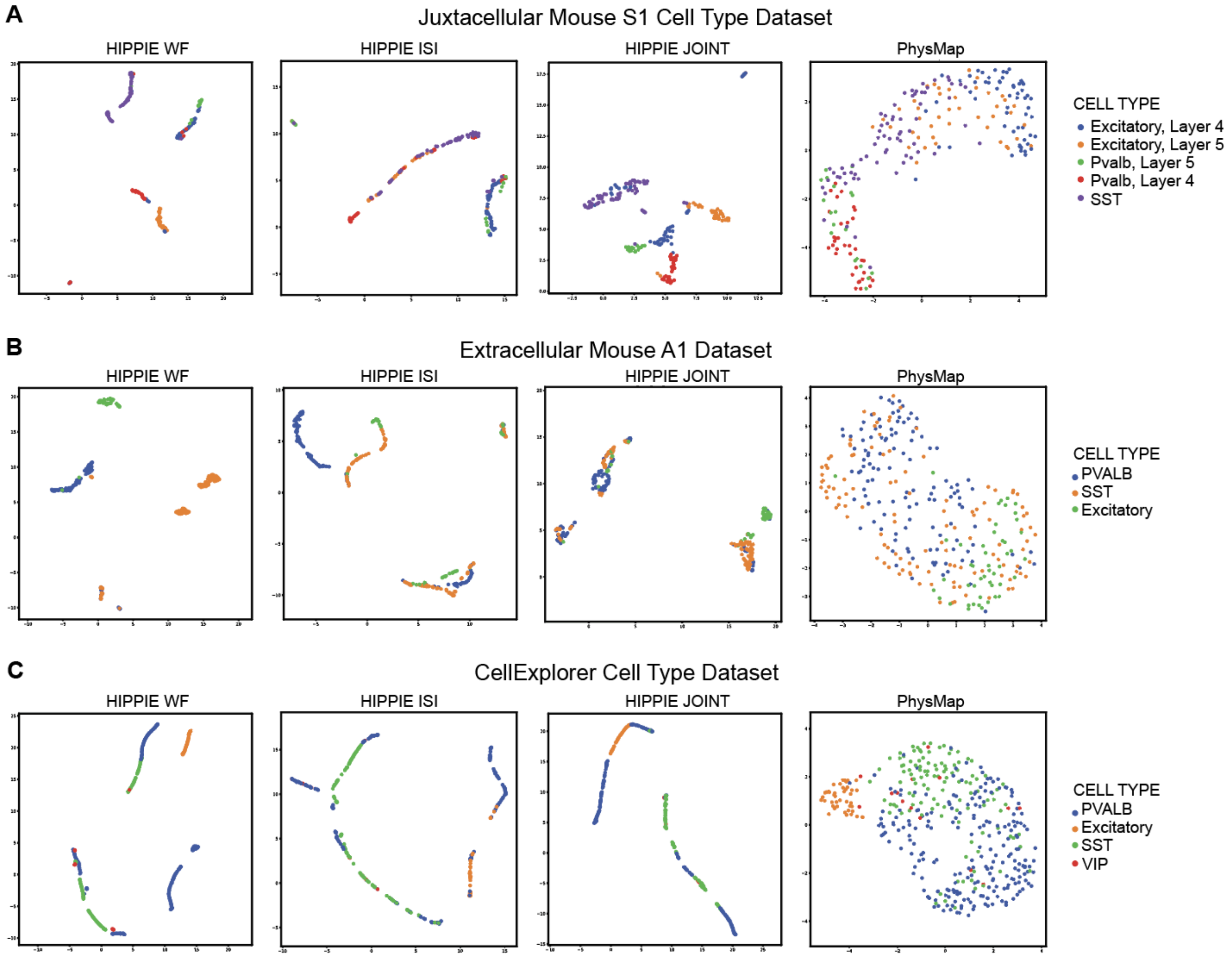


**Supplemental Figure 2**: **UMAP plots for the different dimensionality reduction methods applied to the benchmarks for cell type prediction.** A) UMAP representations for the latent space obtained from HIPPIE (Average Waveform, ISI, Joint) and PhysMap for the Juxtacellular Mouse S1 Cell Type dataset. B) UMAP representations for the latent space obtained from HIPPIE (Average Waveform, ISI, Joint) and PhysMap for the Extracellular Mouse A1 dataset. C) UMAP representations for the latent space obtained from HIPPIE (Average Waveform, ISI, Joint) and PhysMap for the CellExplorer Cell Type dataset.

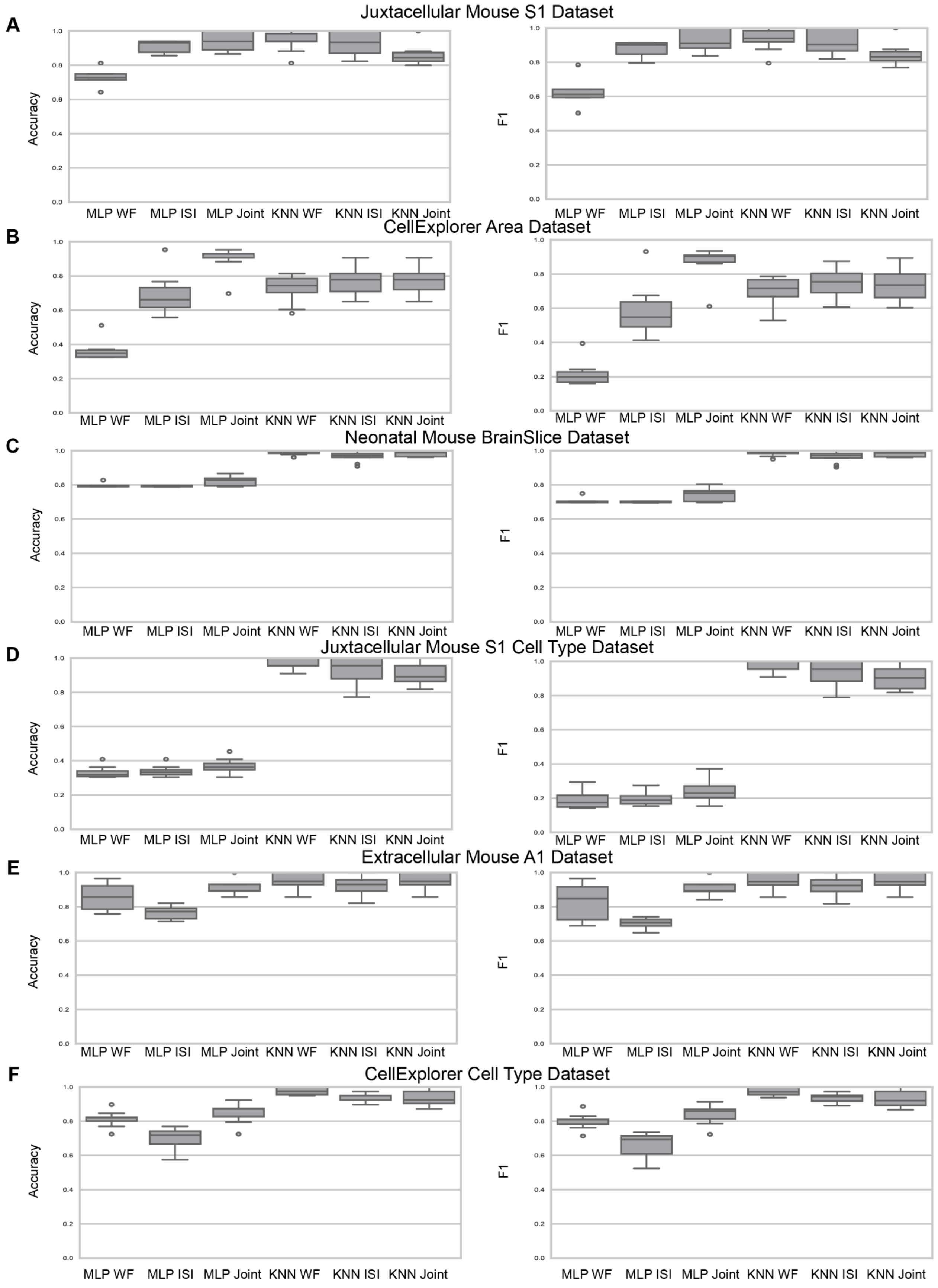
A
Juxtacellular Mouse S1 Dataset
Accuracy
F1
MLP WF
MLP ISI
MLP Joint
KNN WF
KNN ISI
KNN Joint
B
CellExplorer Area Dataset
C
Neonatal Mouse BrainSlice Dataset
D
Juxtacellular Mouse S1 Cell Type Dataset
E
Extracellular Mouse A1 Dataset
F
CellExplorer Cell Type Dataset
0.0
0.2
0.4
0.6
0.8
1.0

**Supplemental Figure 3. K-Nearest Neighbors swap with Multilayer perceptron probe (MLP).** A) Boxplots showcasing the performance of the multilayer perceptron probe on the Juxtacellular Mouse Dataset. B) Boxplots showcasing the performance of the multilayer perceptron probe on the CellExplorer Area dataset. C) Boxplots showcasing the performance of the multilayer perceptron probe on the Mouse Slice Dataset. D) Boxplots showcasing the performance of the multilayer perceptron probe on the Juxtacellular Mouse S1 Cell Type Dataset. E) Boxplots showcasing the performance of the multilayer perceptron probe on the Extracellular Mouse A1 Dataset. F) Boxplots showcasing the performance of the multilayer perceptron probe on the CellExplorer Cell Type Dataset

**Supplemental Table 1. Juxtacellular Mouse S1 Dataset benchmark results.**

| file | accuracy_mean | f1_mean | accuracy_std | f1_std |
|---|---|---|---|---|
| Hippie_wf | 0.9397 | 0.9341 | 0.05811 | 0.06353 |
| Hippie_isi | 0.9309 | 0.9217 | 0.06778 | 0.07153 |
| Hippie_joint | 0.9123 | 0.9080 | 0.06815 | 0.07547 |
| physmap | 0.6876 | 0.6191 | 0.08721 | 0.12234 |
| Raw_isi | 0.6775 | 0.6675 | 0.14774 | 0.14252 |
| Raw_Combined | 0.6502 | 0.6275 | 0.13203 | 0.11886 |
| Raw_WF | 0.6502 | 0.6275 | 0.13203 | 0.11886 |
| PCA_WF | 0.6488 | 0.6000 | 0.07284 | 0.08755 |
| PCA_ISI | 0.6149 | 0.5895 | 0.11013 | 0.10594 |

**Supplemental Table 2. CellExplorer Area Dataset benchmark results**

| file | accuracy_mean | f1_mean | accuracy_std | f1_std |
|---|---|---|---|---|
| hippie_isi | 0.9220 | 0.9196 | 0.05801 | 0.05952 |
| hippie_joint | 0.9504 | 0.9499 | 0.05100 | 0.05128 |
| hippie_wf | 0.9504 | 0.9499 | 0.05100 | 0.05128 |

| | | | | |
|---|---|---|---|---|
| PCA_WF | 0.5587 | 0.4906 | 0.02125 | 0.02084 |
| physmap | 0.5296 | 0.4761 | 0.03732 | 0.03811 |
| PCA_ISI | 0.4945 | 0.4392 | 0.06445 | 0.07370 |
| Raw_WF | 0.4567 | 0.4999 | 0.18201 | 0.21659 |
| Raw_Combined | 0.4558 | 0.4890 | 0.19184 | 0.22508 |
| Raw_isi | 0.3814 | 0.4062 | 0.21628 | 0.22942 |

**Supplemental Table 3. Mouse slice P0 and P3 benchmark results**

| file | accuracy_mean | f1_mean | accuracy_std | f1_std |
|---|---|---|---|---|
| Raw_WF | 0.9705 | 0.9701 | 0.01654 | 0.01691 |
| Raw_isi | 0.9146 | 0.9122 | 0.02013 | 0.02061 |
| Raw_Combined | 0.9711 | 0.9707 | 0.01731 | 0.01768 |
| physmap | 0.9213 | 0.9194 | 0.01685 | 0.01763 |
| PCA_WF | 0.9315 | 0.9244 | 0.01724 | 0.01889 |
| PCA_ISI | 0.8822 | 0.8657 | 0.01439 | 0.01989 |
| Hippie_isi | 0.9662 | 0.9644 | 0.03031 | 0.03251 |
| Hippie_joint | 0.9794 | 0.9790 | 0.01383 | 0.01381 |

**Supplemental Table 4. Juxtacellular Mouse S1 Dataset benchmark results**

| file | accuracy_mean | f1_mean | accuracy_std | f1_std |
|---|---|---|---|---|
| Hippie_isi | 0.9330 | 0.9316 | 0.07738 | 0.07698 |
| Hippie_joint | 0.9065 | 0.9053 | 0.06802 | 0.07024 |
| Hippie_wf | 0.9642 | 0.9642 | 0.03515 | 0.03526 |
| PCA_ISI | 0.4557 | 0.4308 | 0.08423 | 0.08970 |
| PCA_WF | 0.6679 | 0.6335 | 0.03516 | 0.03813 |
| Raw_Combined | 0.7059 | 0.7399 | 0.16507 | 0.17013 |
| Raw_WF | 0.7161 | 0.7545 | 0.16022 | 0.16846 |
| Raw_isi | 0.5808 | 0.6573 | 0.12333 | 0.12562 |

| | | | | |
|---|---|---|---|---|
| physmap | 0.6569 | 0.6070 | 0.06004 | 0.06242 |

**Supplemental Table 5. Extracellular Mouse A1 Dataset benchmark results**

| file | accuracy_mean | f1_mean | accuracy_std | f1_std |
|---|---|---|---|---|
| Hippie_isi | 0.9220 | 0.9196 | 0.05801 | 0.05951 |
| Hippie_joint | 0.9504 | 0.9499 | 0.05100 | 0.05128 |
| Hippie_wf | 0.9504 | 0.9499 | 0.05100 | 0.05128 |
| Raw_isi | 0.6416 | 0.7197 | 0.12812 | 0.11336 |
| physmap | 0.6236 | 0.6128 | 0.08052 | 0.08432 |
| PCA_ISI | 0.6089 | 0.6118 | 0.10253 | 0.10479 |
| Raw_Combined | 0.5808 | 0.6720 | 0.11309 | 0.13034 |
| PCA_WF | 0.5751 | 0.5672 | 0.05393 | 0.05822 |
| Raw_WF | 0.5702 | 0.6564 | 0.11289 | 0.13403 |

**Supplemental Table 6. CellExplorer Cell Type benchmark results**

| file | accuracy_mean | f1_mean | accuracy_std | f1_std |
|---|---|---|---|---|
| Hippie_isi | 0.9413 | 0.9375 | 0.02442 | 0.02749 |
| Hippie_joint | 0.9364 | 0.9316 | 0.04821 | 0.05069 |
| Hippie_wf | 0.9771 | 0.9747 | 0.02244 | 0.02448 |
| Raw_WF | 0.8090 | 0.8346 | 0.15156 | 0.13829 |
| Raw_Combined | 0.8053 | 0.8264 | 0.15923 | 0.14292 |
| PCA_WF | 0.8019 | 0.7838 | 0.05598 | 0.05498 |
| physmap | 0.7781 | 0.7552 | 0.05371 | 0.05584 |
| Raw_isi | 0.7234 | 0.7457 | 0.21607 | 0.19906 |
| PCA_ISI | 0.6863 | 0.6524 | 0.05872 | 0.05987 |

**Supplemental Table 7.** **MLP probe results for all datasets**

| Modality | Mean Accuracy | Mean Macro-F1 | Accuracy standard deviation | Macro-F1 Standard deviation | Dataset |
|---|---|---|---|---|---|
| Hippie_isi | 0.9907 | 0.9881 | 0.01626 | 0.01995 | CellExplorer Area |
| Hippie_joint_mlp | 0.90233 | 0.87106 | 0.07500 | 0.09470 | CellExplorer Area |
| Hippie_joint | 0.7395 | 0.7111 | 0.10389 | 0.11775 | CellExplorer Area |
| Hippie_wf | 0.7395 | 0.7111 | 0.10389 | 0.11775 | CellExplorer Area |
| Hippie_isi_mlp | 0.69070 | 0.58289 | 0.11236 | 0.14659 | CellExplorer Area |
| PCA_WF | 0.5587 | 0.4906 | 0.02125 | 0.02084 | CellExplorer Area |
| physmap | 0.5296 | 0.4761 | 0.03732 | 0.03811 | CellExplorer Area |
| PCA_ISI | 0.4945 | 0.4392 | 0.06445 | 0.07370 | CellExplorer Area |
| Raw_WF | 0.4567 | 0.4999 | 0.18201 | 0.21659 | CellExplorer Area |
| Raw_Combined | 0.4558 | 0.4890 | 0.19184 | 0.22508 | CellExplorer Area |
| Raw_isi | 0.3814 | 0.4062 | 0.21628 | 0.22942 | CellExplorer Area |
| Hippie_wf_mlp | 0.36047 | 0.21516 | 0.05617 | 0.06952 | CellExplorer Area |
| Hippie_joint | 0.9821 | 0.9793 | 0.02714 | 0.03051 | CellExplorer Cell Type |
| Hippie_wf | 0.9821 | 0.9793 | 0.02714 | 0.03051 | CellExplorer Cell Type |
| Hippie_isi | 0.9667 | 0.9597 | 0.04018 | 0.04987 | CellExplorer Cell Type |

| | | | | | |
|---|---|---|---|---|---|
| Hippie_joint_mlp | 0.84974 | 0.84253 | 0.05695 | 0.05703 | CellExplorer Cell Type |
| Hippie_wf_mlp | 0.81397 | 0.80162 | 0.04543 | 0.04503 | CellExplorer Cell Type |
| Raw_WF | 0.8090 | 0.8346 | 0.15156 | 0.13829 | CellExplorer Cell Type |
| Raw_Combined | 0.8053 | 0.8264 | 0.15923 | 0.14292 | CellExplorer Cell Type |
| PCA_WF | 0.8019 | 0.7838 | 0.05598 | 0.05498 | CellExplorer Cell Type |
| physmap | 0.7781 | 0.7552 | 0.05371 | 0.05584 | CellExplorer Cell Type |
| Raw_isi | 0.7234 | 0.7457 | 0.21607 | 0.19906 | CellExplorer Cell Type |
| Hippie_isi_mlp | 0.69917 | 0.65787 | 0.06180 | 0.07705 | CellExplorer Cell Type |
| PCA_ISI | 0.6863 | 0.6524 | 0.05872 | 0.05987 | CellExplorer Cell Type |
| Hippie_joint | 0.9857 | 0.9856 | 0.02497 | 0.02520 | Extracellular Mouse A1 Dataset |
| Hippie_wf | 0.9857 | 0.9856 | 0.02497 | 0.02520 | Extracellular Mouse A1 Dataset |
| Hippie_joint_mlp | 0.90825 | 0.90547 | 0.04223 | 0.04530 | Extracellular Mouse A1 Dataset |
| Hippie_isi | 0.8983 | 0.8969 | 0.07207 | 0.07161 | Extracellular Mouse A1 Dataset |
| Hippie_wf_mlp | 0.85887 | 0.82921 | 0.08030 | 0.11077 | Extracellular Mouse A1 Dataset |
| Hippie_isi_mlp | 0.76502 | 0.70431 | 0.03621 | 0.02827 | Extracellular Mouse A1 Dataset |

| | | | | | |
|---|---|---|---|---|---|
| Raw_isi | 0.6416 | 0.7197 | 0.12812 | 0.11336 | Extracellular Mouse A1 Dataset |
| physmap | 0.6236 | 0.6128 | 0.08052 | 0.08432 | Extracellular Mouse A1 Dataset |
| PCA_ISI | 0.6089 | 0.6118 | 0.10253 | 0.10479 | Extracellular Mouse A1 Dataset |
| Raw_Combined | 0.5808 | 0.6720 | 0.11309 | 0.13034 | Extracellular Mouse A1 Dataset |
| PCA_WF | 0.5751 | 0.5672 | 0.05393 | 0.05822 | Extracellular Mouse A1 Dataset |
| Raw_WF | 0.5702 | 0.6564 | 0.11289 | 0.13403 | Extracellular Mouse A1 Dataset |
| Hippie_isi | 0.9765 | 0.9757 | 0.04113 | 0.04291 | Juxtacellular Mouse S1 Area |
| Hippie_joint_mlp | 0.94203 | 0.93007 | 0.05715 | 0.06410 | Juxtacellular Mouse S1 Area |
| Hippie_joint | 0.9389 | 0.9315 | 0.05012 | 0.05746 | Juxtacellular Mouse S1 Area |
| Hippie_wf | 0.9389 | 0.9315 | 0.05012 | 0.05746 | Juxtacellular Mouse S1 Area |
| Hippie_isi_mlp | 0.91166 | 0.87726 | 0.03457 | 0.04303 | Juxtacellular Mouse S1 Area |
| Hippie_wf_mlp | 0.72313 | 0.61348 | 0.05081 | 0.07953 | Juxtacellular Mouse S1 Area |
| physmap | 0.6876 | 0.6191 | 0.08721 | 0.12234 | Juxtacellular Mouse S1 Area |
| Raw_isi | 0.6775 | 0.6675 | 0.14774 | 0.14252 | Juxtacellular Mouse S1 Area |
| Raw_Combined | 0.6502 | 0.6275 | 0.13203 | 0.11886 | Juxtacellular Mouse S1 Area |

| | | | | | |
|---|---|---|---|---|---|
| Raw_WF | 0.6502 | 0.6275 | 0.13203 | 0.11886 | Juxtacellular Mouse S1 Area |
| PCA_WF | 0.6488 | 0.6000 | 0.07284 | 0.08755 | Juxtacellular Mouse S1 Area |
| PCA_ISI | 0.6149 | 0.5895 | 0.11013 | 0.10594 | Juxtacellular Mouse S1 Area |
| Hippie_joint | 0.9237 | 0.9186 | 0.09550 | 0.09903 | Juxtacellular Mouse S1 Cell Type |
| Hippie_wf | 0.9237 | 0.9186 | 0.09550 | 0.09903 | Juxtacellular Mouse S1 Cell Type |
| Hippie_isi | 0.9105 | 0.9074 | 0.06416 | 0.06866 | Juxtacellular Mouse S1 Cell Type |
| Raw_WF | 0.7161 | 0.7545 | 0.16022 | 0.16846 | Juxtacellular Mouse S1 Cell Type |
| Raw_Combined | 0.7059 | 0.7399 | 0.16507 | 0.17013 | Juxtacellular Mouse S1 Cell Type |
| PCA_WF | 0.6679 | 0.6335 | 0.03516 | 0.03813 | Juxtacellular Mouse S1 Cell Type |
| physmap | 0.6569 | 0.6070 | 0.06004 | 0.06242 | Juxtacellular Mouse S1 Cell Type |
| Raw_isi | 0.5808 | 0.6573 | 0.12333 | 0.12562 | Juxtacellular Mouse S1 Cell Type |
| PCA_ISI | 0.4557 | 0.4308 | 0.08423 | 0.08970 | Juxtacellular Mouse S1 Cell Type |
| Hippie_joint_mlp | 0.36640 | 0.23858 | 0.04359 | 0.06245 | Juxtacellular Mouse S1 Cell Type |
| Hippie_isi_mlp | 0.33933 | 0.19444 | 0.03107 | 0.03834 | Juxtacellular Mouse S1 Cell Type |
| Hippie_wf_mlp | 0.33063 | 0.18866 | 0.03372 | 0.05089 | Juxtacellular Mouse S1 Cell Type |

| | | | | | |
|---|---|---|---|---|---|
| Hippie_isi | 0.9950 | 0.9948 | 0.00616 | 0.00645 | Mouse slice P0 and P3 |
| Hippie_joint | 0.9878 | 0.9865 | 0.01499 | 0.01838 | Mouse slice P0 and P3 |
| Hippie_wf | 0.9878 | 0.9865 | 0.01499 | 0.01838 | Mouse slice P0 and P3 |
| Raw_Combined | 0.9711 | 0.9707 | 0.01731 | 0.01768 | Mouse slice P0 and P3 |
| Raw_WF | 0.9705 | 0.9701 | 0.01654 | 0.01691 | Mouse slice P0 and P3 |
| PCA_WF | 0.9315 | 0.9244 | 0.01724 | 0.01889 | Mouse slice P0 and P3 |
| phys map | 0.9213 | 0.9194 | 0.01685 | 0.01763 | Mouse slice P0 and P3 |
| Raw_isi | 0.9146 | 0.9122 | 0.02013 | 0.02061 | Mouse slice P0 and P3 |
| PCA_ISI | 0.8822 | 0.8657 | 0.01439 | 0.01989 | Mouse slice P0 and P3 |
| Hippie_joint_mlp | 0.82094 | 0.74097 | 0.02689 | 0.03809 | Mouse slice P0 and P3 |
| Hippie_wf_mlp | 0.79579 | 0.70534 | 0.01145 | 0.01590 | Mouse slice P0 and P3 |
| Hippie_isi_mlp | 0.79247 | 0.70073 | 0.00285 | 0.00392 | Mouse slice P0 and P3 |
